# Real time portable genome sequencing for global food security

**DOI:** 10.1101/314526

**Authors:** Laura M. Boykin, Ammar Ghalab, Bruno Rossitto De Marchi, Anders Savill, James M. Wainaina, Tonny Kinene, Stephen Lamb, Myriam Rodrigues, Monica A. Kehoe, Joseph Ndunguru, Fred Tairo, Peter Sseruwagi, Charles Kayuki, Deogratius Mark, Joel Erasto, Hilda Bachwenkizi, Titus Alicai, Geoffrey Okao-Okuja, Phillip Abidrabo, John Francis Osingada, Jimmy Akono, Elijah Ateka, Brenda Muga, Samuel Kiarie

**Affiliations:** School of Molecular Sciences and Australian Research Council Centre of Excellence in Plant Energy Biology, University of Western Australia, Crawley, Perth, WA 6009 Australia; São Paulo State University (UNESP), School of Agriculture, Dept. of Plant Protection, CEP 18610-307, Botucatu (SP), Brazil; Crop Protection Branch, Department of Agriculture and Food Western Australia, South Perth, WA 6151, Australia; Mikocheni Agricultural Research Institute (MARI), P.O. Box 6226, Dar es Salaam, Tanzania; National Crops Resources Research Institute (NaCRRI), P.O. Box 7084, Kampala, Uganda; Jomo Kenyatta University of Agriculture and Technology (JKUAT), P.O. Box 62000 - 00200, Nairobi, Kenya

**Author notes:** Author for correspondence: Laura Boykin.

## Abstract

The United Nations has listed Zero Hunger as one of the 17 global sustainable development goals to end extreme poverty by 2030. Plant viruses are a major constraint to crop production globally causing an estimated $30 billion in damage ^1^ leaving millions of people food insecure ^2^. In Africa, agriculture employs up to 50% of the workforce, yet only contributes 15% to the GDP on average ^3^, suggesting that there is low productivity and limited value addition. This can be addressed through continued innovation in the fields of science and technology as suggested in the Science Agenda for Agriculture in Africa (S3A) ^4^. Sustainable management of plant viruses and their associated vectors must include efficient diagnostics for surveillance, detection and identification to inform disease management, including the development and strategic deployment of virus resistant varieties. To date, researchers have been utilizing conventional methods such as; PCR, qPCR, high throughput sequencing (RNA-Seq, DNA-Seq) and Sanger sequencing for pathogen identification. However, these methods are both costly and time consuming, delaying timely control actions. The emergence of new tools for real-time diagnostics, such as the Oxford Nanopore MinION, have recently proven useful for early detection of Ebola ^6^ and Zika ^7,8^, even in low resourced laboratories. For the first time globally, the MinION portable pocket DNA sequencer was used to sequence whole plant virus genomes. We used this technology to identify the begomoviruses causing the devastating CMD which is ravaging smallholder farmers’ crops in sub-Saharan Africa. Cassava, a carbohydrate crop from which tapioca originates, is a major source of calories for over eight hundred (800) million people worldwide. With this technology, farmers struggling with diseased crops can take immediate, restorative action to improve their livelihoods based on information about the health of their plants, generated using a portable, real-time DNA sequencing device.

## Main

The portable DNA technology has great potential to reduce the risk of community crop failure and help improve livelihoods of millions of people, especially in low resourced communities. Plant diseases are a major cause of low crop productivity and viruses such as Tobacco mosaic virus, Tomato mosaic, Tomato spotted wilt, Potato leaf roll, Potato virus X and Y in Potato, Papaya mosaic, Citrus tristeza, Chilli leaf curl, and Banana bunchy top have been implicated. In particular, cassava viruses are among the world’s greatest risk to food insecurity. Losses caused by cassava mosaic disease (CMD) and cassava brown streak disease are estimated at US $2-3 billion annually ^5^. We therefore visited smallholder farmers in Tanzania, Uganda and Kenya (Table 1) who are suffering yield shortages due to cassava virus infections. We utilized the MinION to test infected material and farmers were informed within 48 hours of the specific strain of the virus that was infecting their cassava, and a resistant cassava variety was deployed. The advantages of adopting this technology far outweigh the challenges (see Table 2). Cassava mosaic begomoviruses were in high enough concentration that reads of whole genomes were obtained without an enrichment step (Table 1). As expected the viral reads increased with the severity of the symptoms observed (Table 1). We detected a dual infection for a leaf sample with the severity score of 5 in Uganda. In addition, one asymptomatic plant in Tanzania had one viral read detected. The shortest time to obtain a viral read was 15 seconds (severity score 5) and the longest was 4h11m15s (severity score 1).

**Table 1.**
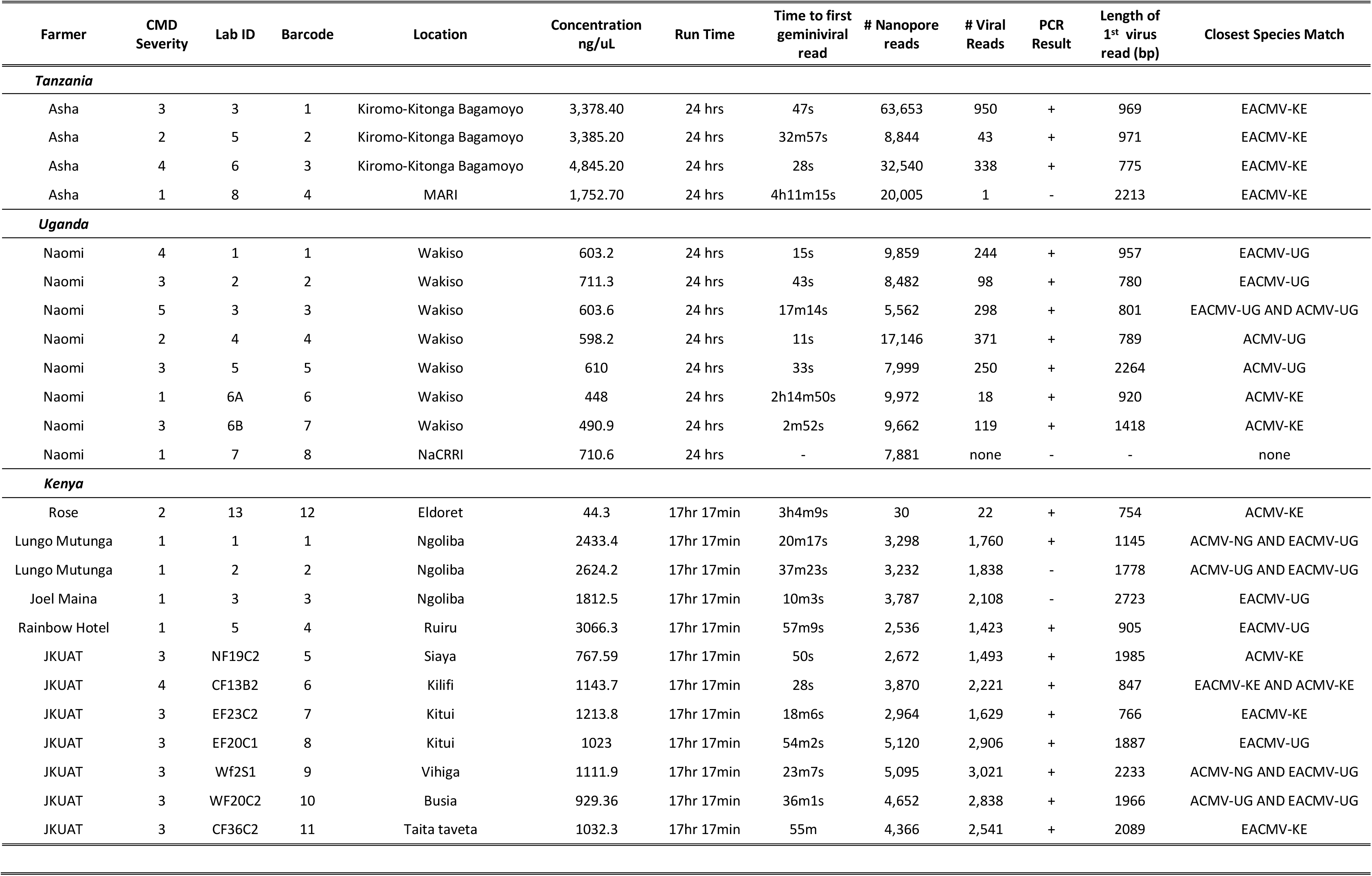
Nanopore results of cassava mosaic begomovirus-infected cassava leaves from Tanzania, Uganda and Kenya. Disease severity described in Legg et al. 2000 where 1 is healthy and 5 shows severe symptoms of the disease including leaf distortion and stunting of the plant (Figure 1).

**Table 2.**
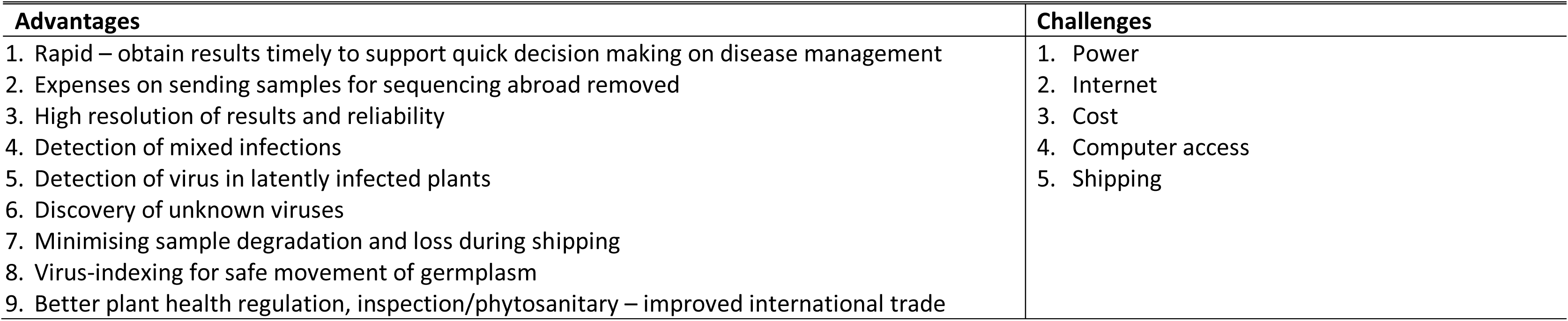
Advantages and challenges of using the MinION portable DNA sequencer in Tanzania, Uganda and Kenya.

| Advantages | Challenges |
| --- | --- |
| 1. Rapid – obtain results timely to support quick decision making on disease management<br>2. Expenses on sending samples for sequencing abroad removed<br>3. High resolution of results and reliability<br>4. Detection of mixed infections<br>5. Detection of virus in latently infected plants<br>6. Discovery of unknown viruses<br>7. Minimising sample degradation and loss during shipping<br>8. Virus-indexing for safe movement of germplasm<br>9. Better plant health regulation, inspection/phytosanitary – improved international trade | 1. Power<br>2. Internet<br>3. Cost<br>4. Computer access<br>5. Shipping |

Additionally, MinION sequencing is superior to traditional methods of PCR identification given its generation of whole genome sequences which enable the identification of the plant virus strain even if it becomes mutated or divergent, as it is not biased using primers that rely on known virus sequences. With regards to cassava, there are three major advantages of this technology. Firstly, improved diagnostics are required and real-time whole genome sequencing will help develop diagnostic primers that are up-to-date. Secondly, this technology will assist with the development of resistant cassava varieties and will allow breeders to immediately test the varieties they are developing against different viral strains. Lastly, it ensures the delivery of the correct healthy uninfected planting material to farmers. In addition, we could detect virus in a plant before it showed symptoms (Table 1). Utilizing traditional PCR methods three samples collected from Asha’s field in Tanzania tested positive for EACMVs and none were positive for ACMV. The asymptomatic sample from MARI tested negative for both ACMV and EACMVs. Eight fresh cassava leaf samples from Uganda were dually infected with ACMV and EACMV-UG using conventional PCR primers for ACMV and EACMV-UG. The primers used in this PCR yield products of 1000bp and 1500bp for ACMV and EACMV-UG respectively. Twelve Kenyan samples were tested and all but two (barcode 2 and 3) were positive using conventional PCR (Table 1). Further studies are needed to verify our results regarding the sensitivity of the protocol for early detection of CMD in cassava, but these results are very promising for ensuring farmers receive clean planting material with early detection of viral infection.

Nanopore sequencing technology has wide applications globally, but in East Africa these include: (a) crop improvement by screening for virus resistant germplasm and genetic diversity during breeding, (b) indexing of cassava planting materials for virus presence or absence to ensure that only clean materials in multiplication fields are distributed to farmers, (c) detection and identification of alternative plant species for cassava-infecting begomoviruses, so that farmers are advised to remove and/or grow crops away from such plants as a management strategy, (d) virus and biodiversity studies.

## Methods

### Sample collection and DNA extraction

In Tanzania, three cassava mosaic disease (CMD) symptomatic cassava leaf samples (Fig. 1, Table 1) were collected from the smallholder cassava farmer Asha Muhammed’s field in Bagamoyo. One more asymptomatic leaf sample was collected at Mikocheni Agricultural Research Institute (MARI), Dar es Salaam. Seven CMD symptomatic plants were collected from Naomi Kutesakwe’s farm in Wakiso district in Uganda. Both Tanzanian and Ugandan samples were collected in September 2017. Twelve samples from Kenya were collected in February 2018 from various sources (Table 1). High quality DNA were isolated using CTAB method ^9^. Each DNA sample (Table 1) was quantified and purity checked using a NanoDrop 2000c UV–vis Spectrophotometer (Thermo Scientific, Wilmington, DE, USA) was used to check the purity and quantity of DNA for each sample and results were recorded in Table 1.

**Figure 1.**
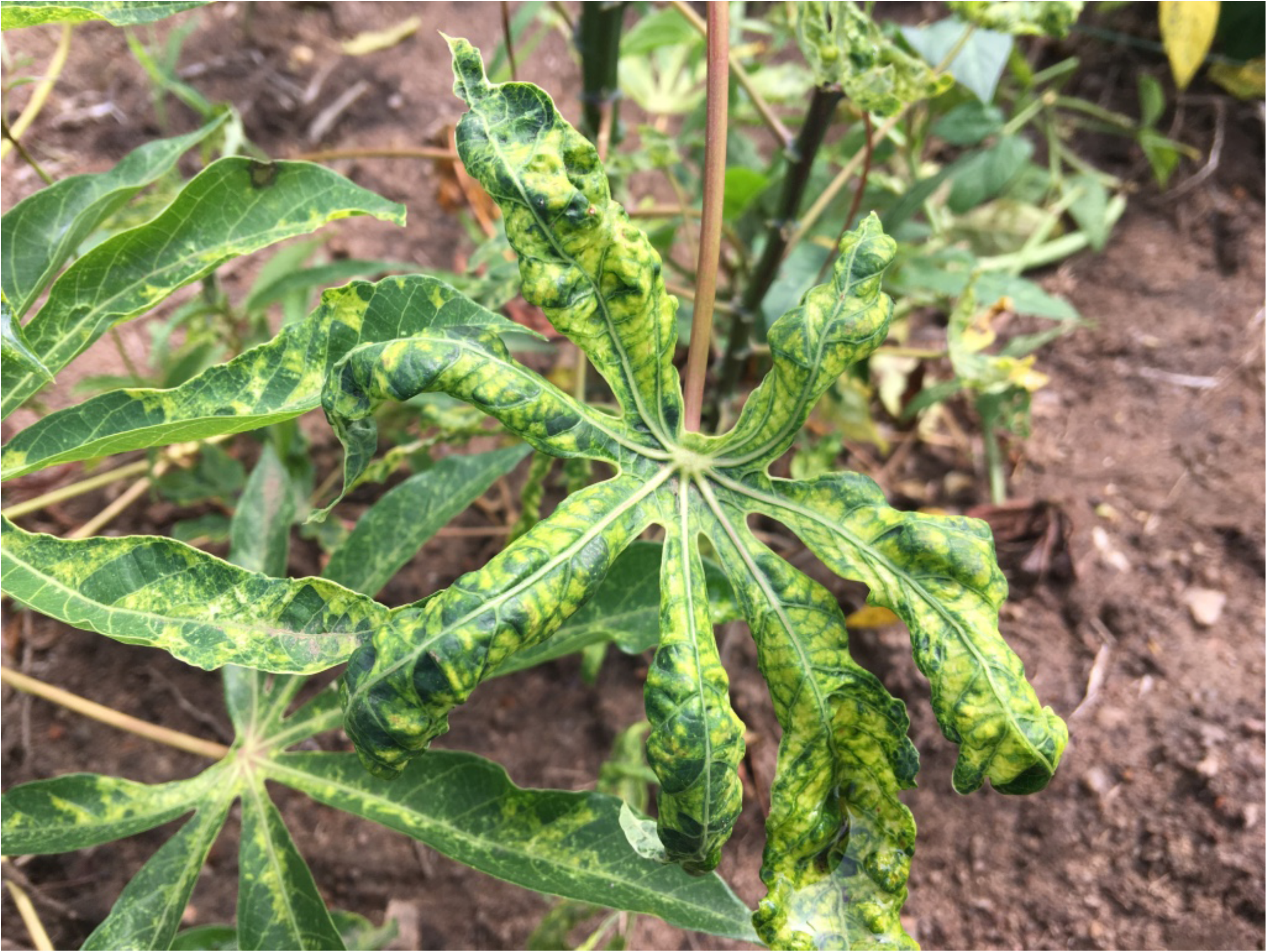
Cassava plant infected with cassava mosaic begomoviruses. Photo credit: Dr. Ndunguru.

### Nanopore library preparation and sequencing

In Tanzania and Uganda, the Rapid Barcoding kit SQK-RBK001 and 9.4.1 flow cells were used to process genomic DNA extracted using a standard CTAB method. We utilized the Rapid Barcoding kit SQK-RBK004 with 9.4.1 flow cells in Kenya. DNA was diluted to 700 ng as specified in the library protocol. The SQK-RBK001 (Sept 2017) and/or the SQK-RBK004 (Feb 2018) protocols were performed as described by the manufacturer. In Tanzania and Uganda, the MinION was run for 24 hours instead of the recommended 48 hours, and in Kenya we had a total run time of approximately 17 hours due to power interruptions.

### Nanopore bioinformatics

In Tanzania and Uganda, Albacore 2.0.2 was used for base calling. In Kenya, Albacore 2.1.10 was used and the scripts were modified to reflect the newest rapid barcoding kit RBK004. Fastq files were imported into Geneious ^10^and a local blast database of all known cassava mosaic begomovirus whole genomes were downloaded from GenBank and a local blast was performed on each of the reads generated using the nanopore device. <u>Results of the local blast were confirmed using blastx on Magnus supercomputer located in Perth, Australia.</u>

### Verification of nanopore results

Traditional PCR was used to verify our nanopore results. In Tanzania and Kenya, two primer pairs: EAB 555F/EAB 555F ^11^and JSP001/JSP002 ^12^, which amplify 556bp and 774bp were used to detect East African cassava mosaic begomoviruses (EACMVs) and African cassava mosaic begomoviruses (ACMVs), respectively. In Uganda, the presence of ACMV and EACMV in each sample was detected using a pair of specific primers for ACMV, ACMV-AL1/F and ACMV-ARO/R and specific for EACMV-UG2, UV-AL1/F and ACMC-CP/R3 ^13^.

## Acknowledgements

Funding for the Kenyan trip was provided by the Crawford Fund. We also thank the participants from the University of Eldoret who assisted in the preparation of libraries for the Kenyan samples.

## Author contributions

Designed the study: LMB, JN, TA, FT, PS, MK, AS, EA. Carried out experiments: LMB, AG, BD, JW, MR, JN, PS, CK, DM, JE, HB, TA, GO, PA, JO, JA, EA, BM and SK. Analysed data: LMB, AS, CK, DM, JE, HB, SL, JN, PS, TA, GO, PA, JO, JA, EA, BM. Contributed to the writing: All authors contributed to the writing of the manuscript.

